# From taxonomic to functional dark diversity: exploring the causes of potential biodiversity and its implications for conservation

**DOI:** 10.1101/2020.03.09.984435

**Authors:** Loïs Morel, Vincent Jung, Simon Chollet, Frédéric Ysnel, Lou Barbe

**Affiliations:** UMR BOREA, MNHN, CNRS, UPMC, IRD, UC, UA, Université de Rennes 1, Campus Beaulieu, 35042, Rennes Cedex, France; UMR 6553 ECOBIO, OSUR, CNRS, Université de Rennes 1, 35042 Rennes, France

**Keywords:** community assembly, ecological restoration, forest temporal continuity, passive rewilding, plants, spiders, taxonomic, functional dark diversity

## Abstract

1. Dark diversity is an emerging and promising concept proposed to estimate the recruitment potential of natural communities and guide conservation and restoration policies. It represents all the species that could be present in a community due to favourable environmental conditions, but are currently lacking. To date, experimental approaches only measured taxonomic dark diversity, mainly based on species coexistence, which relies partly on neutral processes. Thus, these approaches may fail to identify the biodiversity which is lacking for deterministic reasons, and can hence hardly bring out suitable restoration methods.
2. Here, we propose a novel method to estimate dark diversity, which is based on more deterministic coexistence: the coexistence of species’ functional features. We adapted the Beals’ co-occurrence index using functional groups, and we estimated functional dark diversity based on coexistence of functional groups. We then made use of functional dark diversity to address a persistent issue of restoration ecology: how does passive rewilding impact the ecological integrity of recovered communities? We compared spontaneous, secondary woodlands with ancient forests, in terms of taxonomic and functional dark diversity of vascular plants and spiders.
3. Our results indicated that functional dark diversity does not equate to taxonomic dark diversity. Considering plants, recent woodlands surprisingly harboured less functional dark diversity than ancient forests, while they had a very similar amount of taxonomic dark diversity. Concerning spiders, recent woodlands harboured a similar amount of functional dark diversity as ancient forests, but more taxonomic dark diversity. Also, the composition of functional dark diversity differed between forest types, shedding light on their past assembly processes and unveiling their potential for conservation and effective restoration.
4. *Synthesis and applications*. Functional dark diversity brings novel perspectives for ecological diagnostic and restoration. Combined to taxonomic dark diversity, it enables to identify easily the deterministic constrains which limit the re-assembly of ecological communities after land-use changes and to predict the realistic, possible establishments of functional features. Here, we showed that spontaneous woodlands can have very similar, sometimes even higher, ecological integrity than that of ancient forests, and hence may be valuable habitats to be conserved from an ecological perspective.

## Introduction

The concept of *dark diversity* has recently been introduced in Ecology by Pärtel, Szava-Kovats, & Zobel (2011), to take into account the potential biodiversity of natural communities. In a given community, the dark diversity represents the diversity of species that are locally absent while they are present in the regional pool and could be present due to favourable environmental conditions (*i.e.* they are present in the habitat-specific species pool, Pärtel et al., 2011). Therefore, dark diversity identifies species that are absent due to dispersal limitation or historical contingencies, but not species that are absent due to recruitment limitation (that should be absent anyway) nor species that the sampling failed to observe (dormant or very rare species, see Pärtel, 2014). Dark diversity places biodiversity into a dynamic perspective: for example, it integrates species with a colonisation credit, which are species likely to be recruited in the future due to favourable environmental conditions or delayed population growth (Jackson & Sax, 2010). Moreover, dark diversity sheds light on assembly processes, for example by determining the extent to which stochastic processes such as dispersal influence assembly (Pärtel, 2014; Pärtel et al., 2011). Consequently, identifying the dark diversity of communities enables to guide the conservation efforts and the restoration strategies, since it helps to determine the taxa that are frequently absent (e.g. see Moeslund et al., 2017), the habitats more or less degraded, their restoration potential, and, conversely, the habitats that are the most complete (*i.e.* with the lowest dark diversity, Lewis et al., 2017) and that could hence be priority targets for conservation. However, to date, the studies evaluating dark diversity are restricted to the taxonomic facet of communities, hence to species’ identities.

Incorporating functional traits of species into direct assessments of dark diversity could bring many novel insights. Functional traits are all the features of species that can either respond to environmental conditions or can impact ecosystem functions, or both (Violle et al., 2007). Basically, these are morphological, physiological or phenological features, for example the life form of a plant or the type of diet of an animal. The tools that are currently available for measuring directly dark diversity do not consider functional traits, overlooking the ecological differences that may exist between or within taxa. Functional traits can obviously be very different among taxa but also within taxa (Prinzing et al., 2008), and in taxa occupying particular environments (Hermant, Hennion, Bartish, Yguel, & Prinzing, 2012). Alternatively, at some trophic level, several species can have similar functional traits and can therefore be redundant in the impact they have on ecosystem functions or in the response they have to disturbances or environmental changes. Consequently, a given taxonomic dark diversity could or could not represent a functional dark diversity (Figure 1b, c, d), which would bring very different information about past assembly processes, the potential outcome of community assembly, as well as the interest for conservation and the capacity of communities to be ecologically restored.

**Figure 1.**
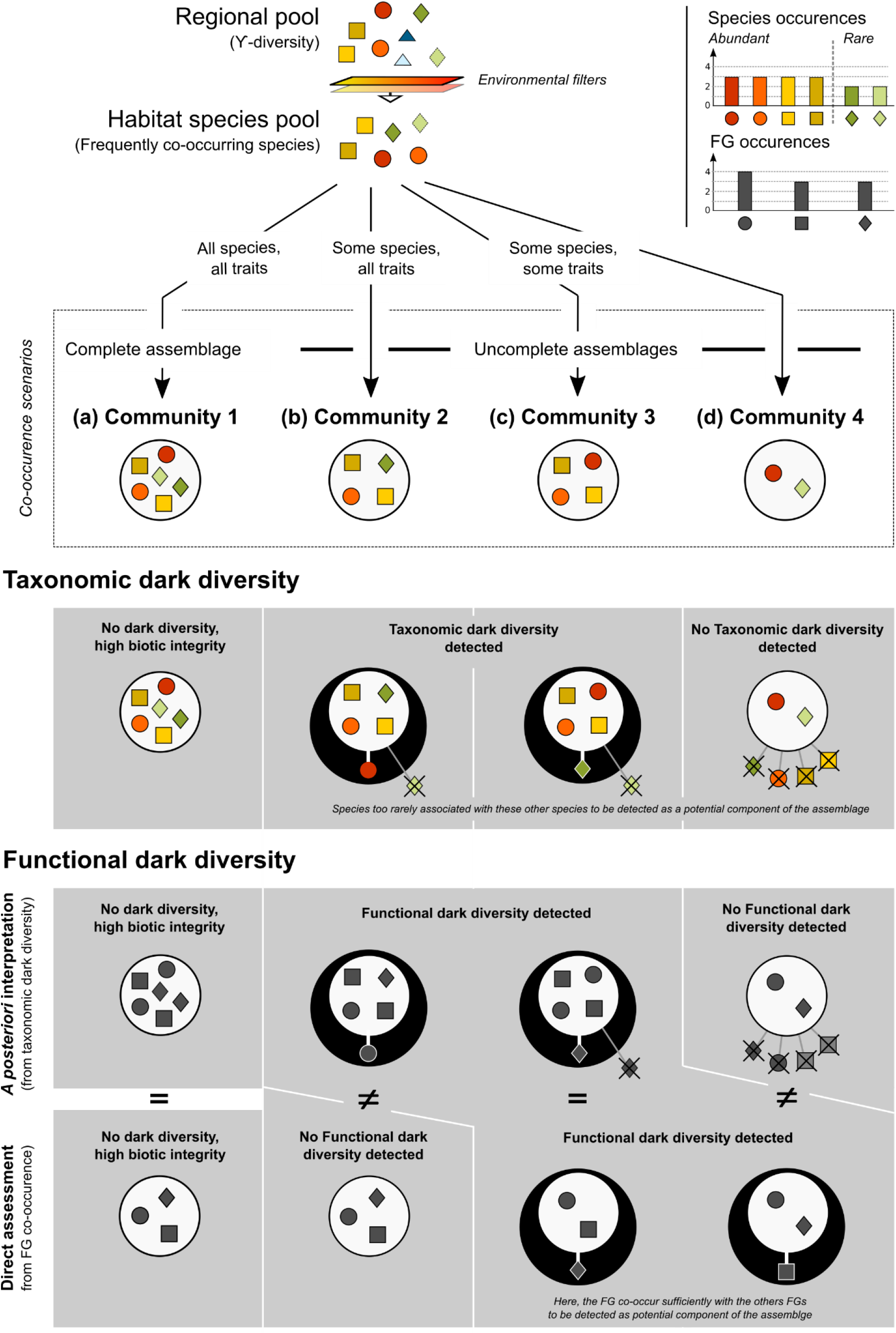
Four distinct co-occurrence scenarios in several communities (a, b, c, d) from a single habitat-specific species pool. The colour of drawing represent a species identity, and its shape represent its functional group. From these local co-occurrences taken together, taxonomic dark diversity and functional dark diversity can be estimated for the four communities. Functional dark diversity estimated *a posteriori* (*i.e.* functional interpretation of taxonomic dark diversity) mirrors taxonomic dark diversity, because it is only the translation into traits of the species which are lacking. Functional dark diversity estimated *a priori* from the co-occurrence of functional groups differs from taxonomic dark diversity and hence from functional dark diversity estimated *a posteriori.* In particular, in the second community (b), a species is lacking, so its functional group seems also lacking. However, this functional group is in fact already present in the community so cannot be recruited. In the fourth community (d), many species are lacking but their co-occurrences with the present species are too rare, so these species are not integrated into the taxonomic dark diversity. However, the co-occurrence of their functional groups is frequent, thus the lacking functional group, despite hosted by rare species, is integrated into the functional dark diversity calculated *a priori*.

While a functional interpretation of taxonomic dark diversity can be made *a posteriori* (*i.e.* what are the traits of the species which are absent?), we think that the calculation of taxonomic dark diversity, which is based in most cases on co-occurrence of species (Lewis, Szava-Kovats, & Pärtel, 2016), can in itself limit the detection of functional patterns. Variations of taxonomic diversity have been shown to often result from non-deterministic processes such as neutral coexistence (Chase & Leibold, 2003), suggesting that taxonomic diversity could be a somewhat unreliable and unpredictable parameter, influenced by stochastic processes. Moreover, taxonomic dark diversity taken alone can potentially underestimate the ecological integrity of a community (*i.e.* the capacity of a community to harbour species composition, diversity and functional organisation similar to those of undisturbed ecosystems in the region) because it does not consider the functional redundancy within taxa and the niche filling within habitats (Figure 1b). Most importantly, modern ecology has shown that, from an ecological perspective, the coexistence of functional features, which can be traits or combinations of traits, is much more informative and relevant than the coexistence of species (Mcgill, Enquist, Weiher, & Westoby, 2006). What coexist are functional features, much more than truly independent species: for plants, for example chamephytes, therophytes and small hemicryptophytes in peatlands, helophytes and hydrophytes in marshes, woody species, shrubs, lianas and small herbs in forests… Consequently, accounting for functional features in the co-occurrence calculation becomes a necessity if we want to correctly infer from the concept of dark diversity the ecological determinants of community assembly and identify the potential functions that can realistically be recruited in natural communities, and thus, take fully appropriate conservation and restoration policies. Last, an assessment of functional dark diversity through coexistence of functional features would also increase the probability of detecting functional features hosted by rare species: the co-occurrences of these species might be too rare for them to be included in the taxonomic dark diversity (so their groups would not be included either with an *a posteriori* interpretation), but the co-occurrences of their groups may be sufficiently frequent for the groups to be included in the functional dark diversity (Figure 1d).

Terrestrial ecosystems currently experience many land-use changes, which raises important questions about their impacts on biodiversity and natural habitats (Newbold et al., 2015). In particular, how communities of ancient forests differ from those of recent woodlands is an old but persistent issue of ecology (Bergès & Dupouey, 2020), which offers an ideal opportunity to make use of dark diversity. Recent woodlands are spontaneous forests resulting from a secondary succession following land abandonment, whereas ancient forests are uninterrupted forests since several centuries (at least 150-400 years in western Europe, Hermy, Honnay, Firbank, Grashof-Bokdam, & Lawesson, 1999). The interruption of forest temporal continuity generally induces two major constraints for the forest re-assembly: recruitment limitation and dispersal limitation, respectively due to past land-uses (e.g. fertilisation or soil disturbance) and spatio-temporal fragmentation of source habitats (Hermy & Verheyen, 2007; Kimberley, Blackburn, Whyatt, Kirby, & Smart, 2013). Consequently, the composition and structure of communities in recent woodlands often differ strongly from those of ancient forests. Notably, recent woodlands often lack specialised, typical plant species of ancient forests, which are characterised by low seed production, low dispersal capacities, and require very precise ecological conditions such as oligotrophic substrates and soils weakly disturbed (Flinn & Vellend, 2005). Animal groups may also be sensitive to the forest temporal continuity, in particular arthropods, which are highly dependent to local habitat conditions (Hofmeister et al., 2019). Among arthropods, spiders might be particularly interesting to survey because they are ubiquitous in all terrestrial ecosystems, and the structure of their community might be gradually reshaped during the successional trajectory (Morel et al., 2019; Oxbrough, Gittings, O’Halloran, Giller, & Smith, 2005). Overall, many aspects of the ecological consequences of the rupture of forest temporal continuity remain to be deepened, for instance the relative importance of dispersal and recruitment limitations on biodiversity recovering, which is highly context-dependant (see Bergès & Dupouey, 2020). Thus, the application of the dark diversity framework should enable to obtain a more realistic vision of the capacities of ecosystems to spontaneously recover biodiversity.

Here, we developed the first method to estimate functional dark diversity, and we applied this method to evaluate how passive rewilding (*i.e.* spontaneous afforestation) may reshape the biocenosis of forest ecosystems. We sampled plant and spider communities, two main understorey taxas which depend on distinct biotic and abiotic conditions, and characterised their spectrum of functional traits. We calculated taxonomic dark diversity using the species co-occurrence method (Lewis et al., 2016) and adapted this method to assess functional dark diversity, using co-occurrence of functional groups, which were identified through multitrait differences. Then, we compared recent woodlands with ancient forests. We tested the following hypotheses: (i) the composition of functional dark diversity is specific to the forest type, illustrating their different ecological constraints, (ii) recent woodlands harbour plant and spider communities with both higher taxonomic and functional dark diversity than ancient forests (*i.e.* restoration is partially effective) and (iii) functional dark diversity does not equate to taxonomic dark diversity.

## Materials and methods

### Study sites

We conducted the study in different forest environments of Western Europe (Brittany, France). We selected 32 plots of mesophilic, oak and beech-dominated mature forests, within sites sharing similar geological substrates (mainly granite rocks and schists), thereby strongly limiting the influence of environmental heterogeneity and stand maturity. These plots were homogeneous management units of around 1 ha and were distributed across 8 sites (ranging from 200 to 4000 ha) within the regional area. We set apart plots of ancient forests from those of recent woodlands by checking the temporal forest continuity thanks to the historical Cassini map layers (year 1790) and the Napoleonic cadastre (year 1847), that is, the two reference documents in France for the historical land-uses (Cateau et al., 2015). We defined ancient forests as sites already forested in the middle of the 18^th^ century (when the overall forested area was at its minimum over the French territory, Cateau et al., 2015) and recent woodlands as forests resulting from farmland abandonment during the 20^th^ century. Therefore, ancient forests have an uninterrupted forest state since at least 230 years and recent woodlands are not older than 120 years. Our dataset included 20 plots in ancient forests (from six different forest sites) and 12 plots in recent woodlands (from two different forest sites). The habitat structure and the ecological conditions were quite similar between recent and ancient forests: there were no differences of canopy cover, basal area and Ellenberg Indicator Values (EIV) for moisture degree (Table S1). But, EIV showed higher pH, nutrient concentration and shading in recent woodlands in comparison to ancient forests (Table S1), which is consistent with previous studies investigating environmental conditions in post-agricultural woodlands (Koerner, Dupouey, Dambrine, & Benoit, 1997).

### Community sampling

We conducted floristic surveys in June-July 2014 and 2015 to sample the understorey plant communities of the selected plots (*i.e.* below 2 meters high and including woody species). We used 50-m^2^ quadrats (10 x 5 m) and we noted all species encountered belonging to the herbaceous and shrub strata. A total of 99 species was recorded.

To sample spider communities, we compiled data from a regional database which included individual sampling conducted within the same 32 plots that we used for the floristic surveys. Sampling was made using a standardised protocol based on 3 pitfall traps spaced 10m apart and located at the centre of the plot. Sampling was conducted between April and June either in 2013, 2014, or 2015 (see Morel et al., 2019 and references for database description and more details on the sampling method). The final dataset included 3615 adult individuals, belonging to 89 species.

### Functional characterisation of species

We selected 9 functional traits from the LEDA database (Kleyer et al., 2008) to measure the functional variability of plant species. These traits relate to the plants’ ecological strategy for resource acquisition, competition, regeneration and dispersal (Table S2). We selected two traits of the leaf economics spectrum (Wright et al., 2004) informing about resource acquisition, resource conservation and competition: the specific leaf area (SLA) and the leaf dry matter content (LDMC). We selected four regenerative traits related to growth and dispersal in space and time (Pérez- Harguindeguy et al., 2013): dispersal syndrome, pollination type, seed dry mass and start of flowering. We also selected two integrative traits informing about the overall ecological strategy of plants: plant height and life form. All of these traits are response-effect traits (Lavorel & Garnier, 2002) since they both respond to environmental conditions and also influence ecosystem functions. Since traits were not overly correlated (all r < 0.60), we kept the 9 selected traits. The dataset comprised 16 missing values, that is, 1.8% of the dataset.

We selected 4 life-history traits available in the literature to characterise the functional variability of spider species (Table S3): body size, guild, phenology and circadian activity. These traits relate to the ecological strategy of spiders, in particular their diet and hunting specialisation, foraging method, the habitats they exploit and their dispersal abilities. They hence represent key features illustrating the assemblages of predator arthropods at local scale (Cardoso, Pekár, Jocqué, & Coddington, 2011).

### Identification of functional groups

Since we aimed at using a co-occurrence index to calculate functional dark diversity, we needed to divide the species pool into discontinuous elements, that is, functional groups. For plants, we divided the species pool into functional groups following the methods of Verheyen, Honnay, Motzkin, Hermy, & Foster (2003), which were used in a similar investigation of recent versus ancient understorey plant communities. This method allows to identify functional groups according to trait correlations and thus select objectively consistent ecological groups. First, we calculated a species-to-species distance matrix with the Gower’s similarity coefficient, since this coefficient can deal with missing values and both quantitative and qualitative data. Then, we used this matrix to cluster the species into functional groups using the Ward’s method (Murtagh & Legendre, 2014). The optimal number of groups was determined graphically from visual screening of the dendrogram (Figure S1). We identified 10 functional groups of plant species: 4 groups of woody plants and 6 groups of herbaceous and graminoid species (Figure S1). We made sure that the selected groups were ecologically relevant, that is, corresponded to subsets that were noticeable on the field. For spiders, we applied the method of functional entities since all traits were categorical, and each unique combination of traits resulted in a distinct group (Mouillot et al., 2014). Thus, we identified 35 functional entities (Table S4). We run the further analyses with these groups for plants and spiders, but note that we also run the analyses with groups defined *a priori*, to ensure that our results were not trivially the reflection of group selection. For plants, we adapted the “biological types” of species recorded in the French flora database (Baseflor; Julve, 1998), which are derived from the classification of Raunkier, and we partitioned the species into 12 groups. For spiders, we used the guilds’ typology developed by Cardoso et al. (2011), and we partitioned the species into 7 groups. With this alternative group selection, we obtained the same results hereafter for both plants and spiders (Figure S2).

### Measuring dark diversity and completeness

First of all, we measured the taxonomic and functional, observed diversity of communities (see Table S5). Then, we calculated dark diversity using the Beals’ co-occurrence index (Beals, 1984), a method considered by Lewis et al. (2016) as one of the most efficient. This method relies on a calculus of co-occurrence that enables to identify the subset of species that have the greatest probabilities to coexist, within the habitat-specific species pool that was defined as our whole dataset. In a given community, the taxonomic dark diversity integrates species that are absent but have the greatest probabilities to coexist with the present species (Figure 2). We calculated taxonomic dark diversity according to this method, using a significance threshold of 0.05, as advised by (Lewis et al., 2016).

**Figure 2.**
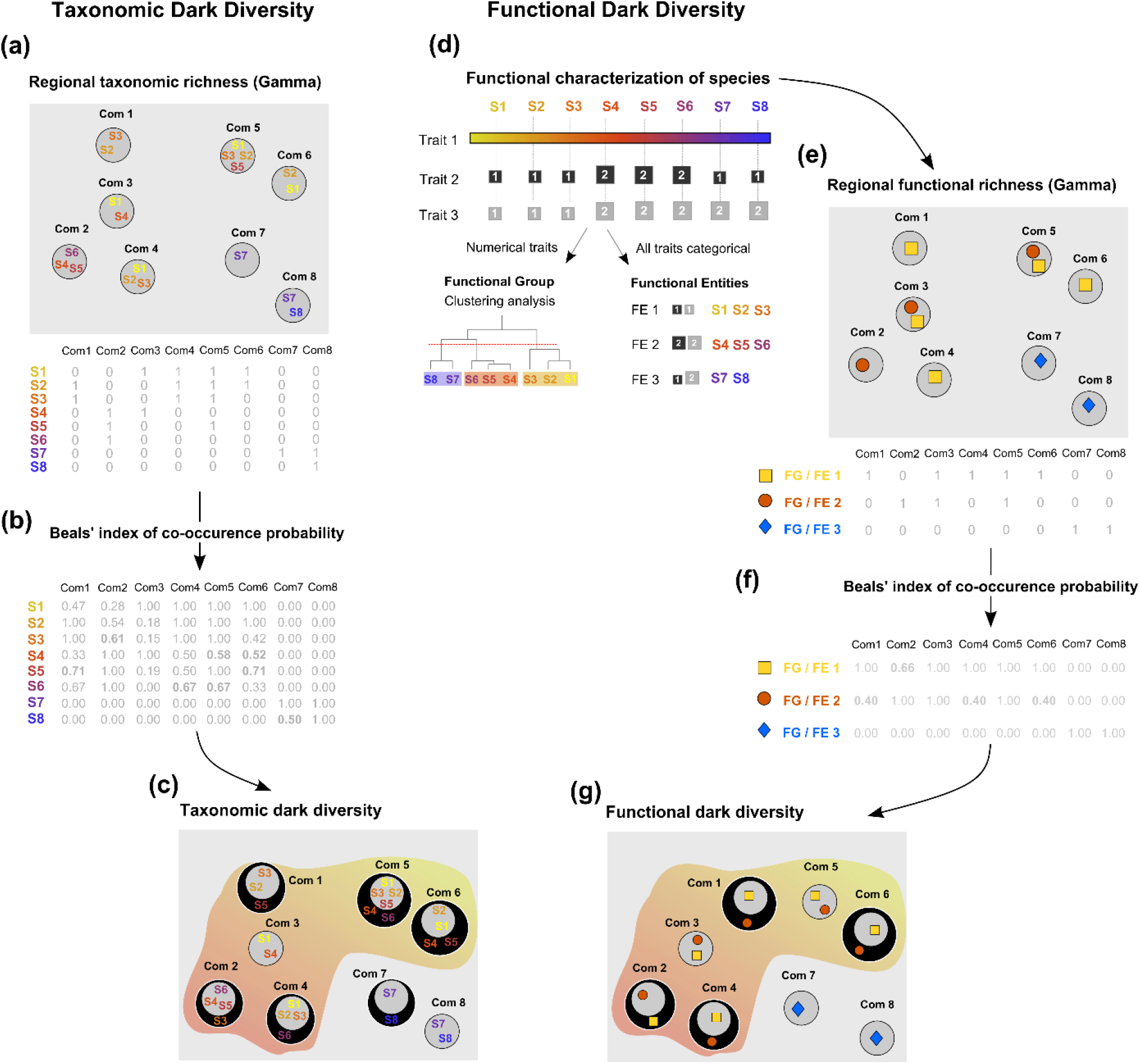
Analysis approach for measuring taxonomic and functional dark diversity. “Com1” means community 1 and “S1” means species 1. (a) From the species’ presence/absence matrix, (b) the Beals’ index estimates the co-occurrence probability of each species in each community. (c) In a given community, a missing species having a high co-occurrence probability in this community will be integrated to the taxonomic dark diversity of this community. The methodological principle for measuring functional dark diversity is identical (e, f, g), but the Beals’ index relies on a matrix of functional groups (or functional entities), preliminary obtained from the functional characterization of species (d). In (c) and (g), the dark diversity of each community is represented by a circle with a black background surrounding the communities with their observed diversity.

Then, we adapted this method to calculate functional dark diversity: instead of using taxonomic co-occurrence, we used functional co-occurrence, that is, the probability of functional groups or functional entities to coexist (Figure 2). The rest of the procedure was identical: we identified in each community the functional groups that were absent while they have an important probability to coexist with the functional groups present in the community. We also calculated a percentage of change between recent and ancient forest for each species and each functional group identified in the dark diversity. Finally, to avoid biased interpretations of the differences in dark diversity due to variations in species richness, we calculated the functional completeness of communities (Pärtel, Szava-Kovats, & Zobel, 2013), that is, their observed diversity relative to their dark diversity. We used the formula: *In* 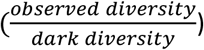. The numerator and denominator were increased by 1 to avoid the limits of division by zero (Helm, Zobel, Moles, Szava-Kovats, & Pärtel, 2015).

### Data analysis

We had a dataset with a nested structure: replicate plots nested into forest plots, nested into forest types. Therefore, we used generalised linear mixed models (GLMMs with Poisson distribution family) for discrete variables (dark diversity) and linear mixed models (LMMs) for continuous variables (completeness) to test differences among forest types (*i.e.* ancient versus recent). We used the forest type as a fixed factor and the hierarchical structure (plots nested within sites) as a random effect, to remove any potential effect of autocorrelation. All analyses were performed using R software (R core team, 2017). The handling of trait matrices and identification of functional groups were done using the package “cluster” and the “species_to_FE” and “FE_metrics” functions (Mouillot et al., 2014). The measures of dark diversity were made with the package “vegan” and the “beals” function, and the script provided by Lewis et al. (2016). Statistical tests were performed thanks to the package “lme4”.

## Results

### Composition of dark diversity in ancient and recent forests

The composition of taxonomic and functional dark diversity strongly differed between both forest types. Only 4 plant species, 4 spider species, 3 plant functional groups and 9 spider functional entities were observed in the dark diversity of both forest types. In the other hand, 10 plant species and two plant functional groups (mesophanerophytes and vernal geophytes) were specific to the dark diversity of ancient forests (Figure 3), whereas 9 plant species and one plant functional group (megaphanerophytes) were specific to the dark diversity of recent forests (Figure 3). Furthermore, 11 spider species and 6 spider functional entities were specific to the dark diversity of ancient forests and 8 spider species and one spider functional entity were specific to the dark diversity of recent forests (Figure 3). Last, we observed that some species were present in the taxonomic dark diversity while their groups were absent in the functional dark diversity, and conversely (Figure 3).

**Figure 3.**
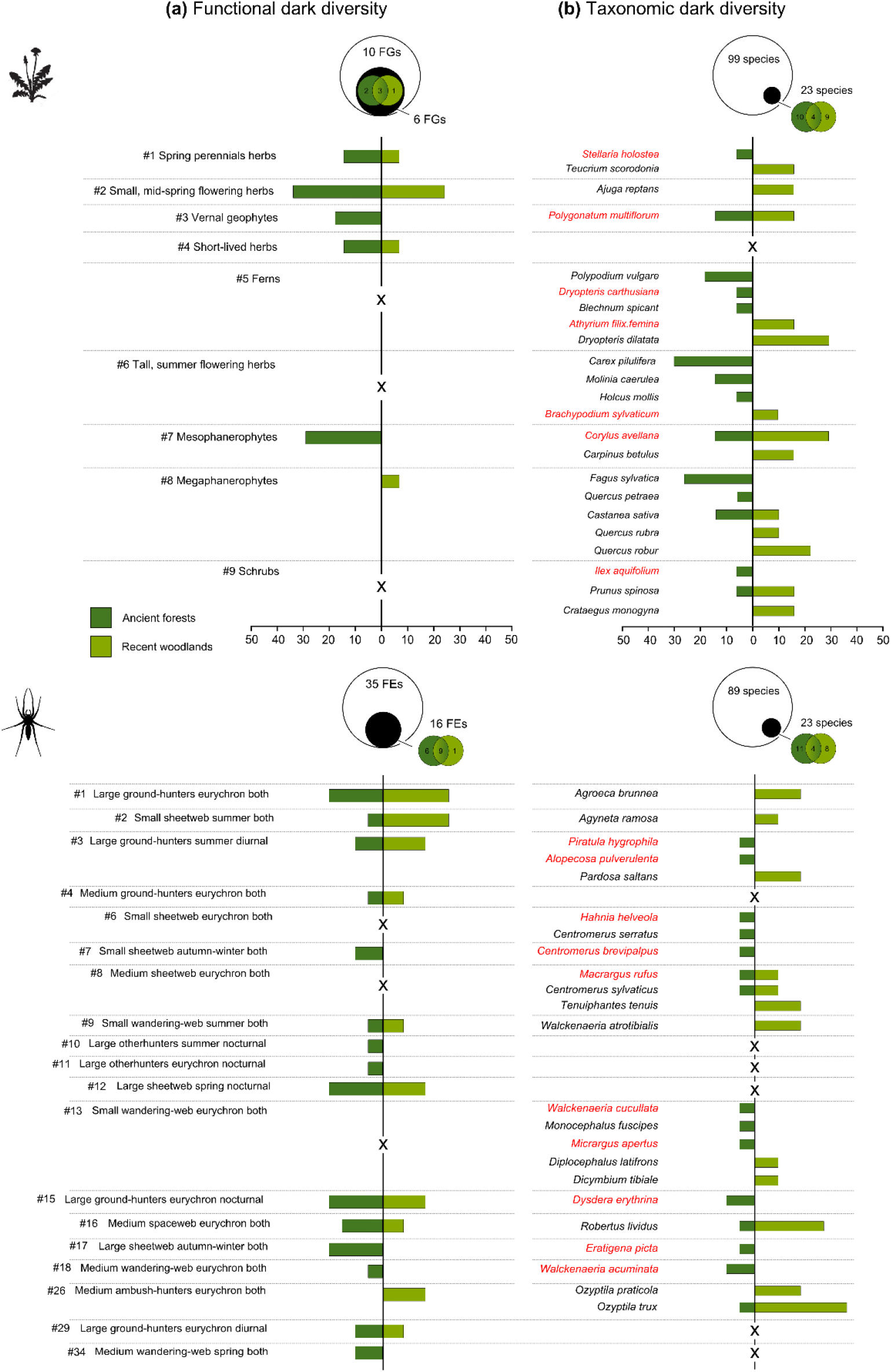
Occurrence frequency in the dark diversity of ancient and recent forests of (a) functional groups (FGs) of plants and functional entities (FEs) of spiders and (b) species of plants and spiders. Only groups, entities and species observed at least once in the dark diversity are presented. Species usually more frequent in ancient forests are in red (from Hermy et al., 1999 and Morel et al., 2019 for plants and spiders, respectively). Black crosses indicate absence in the dark diversity. See Figure S1 and Table S4 for more details on functional groups and entities. Also, see Table S5 for compare species and FG/FE occurrence frequency in observed diversity.

### Taxonomic and functional dark diversity in ancient and recent forests

In total, 23 plant species (23% of the species pool) and 23 spider species (26% of the species pool) were recorded at least once in the taxonomic dark diversity (Figure 3). Also, 6 functional groups of plants (60% of the pool of functional groups) and 16 functional entities of spiders (46% of the pool of functional entities) were recorded at least once in the functional dark diversity (Figure 3).

Overall, we found differences in taxonomic and functional dark diversity between ancient and recent forests (Figure 4a). We found these differences were opposite for taxonomic and functional dark diversity, both in sign and magnitude. For plants, there was similar taxonomic dark diversity in ancient and recent forests (2.3 ± 2.5 vs. 1.8 ± 1.4, p>0.05, Wald’s test) but more functional dark diversity in ancient forests (1.2 ± 1.1 vs. 0.5 ± 0.8, p<0.05, Wald’s test). For spiders, there were more taxonomic dark diversity in recent forests than in ancient ones (1.8 ± 1.7 vs. 0.9 ± 0.9, p<0.05, Wald’s test), but similar functional dark diversity (1.5 ± 1.2 vs. 1.7 ± 1.7, p> 0.05, Wald’s test). Last, the completeness of recent woodlands was higher than that of ancient forests concerning plant functional group, and similar concerning spider functional entities (Figure 4b).

**Figure 4.**
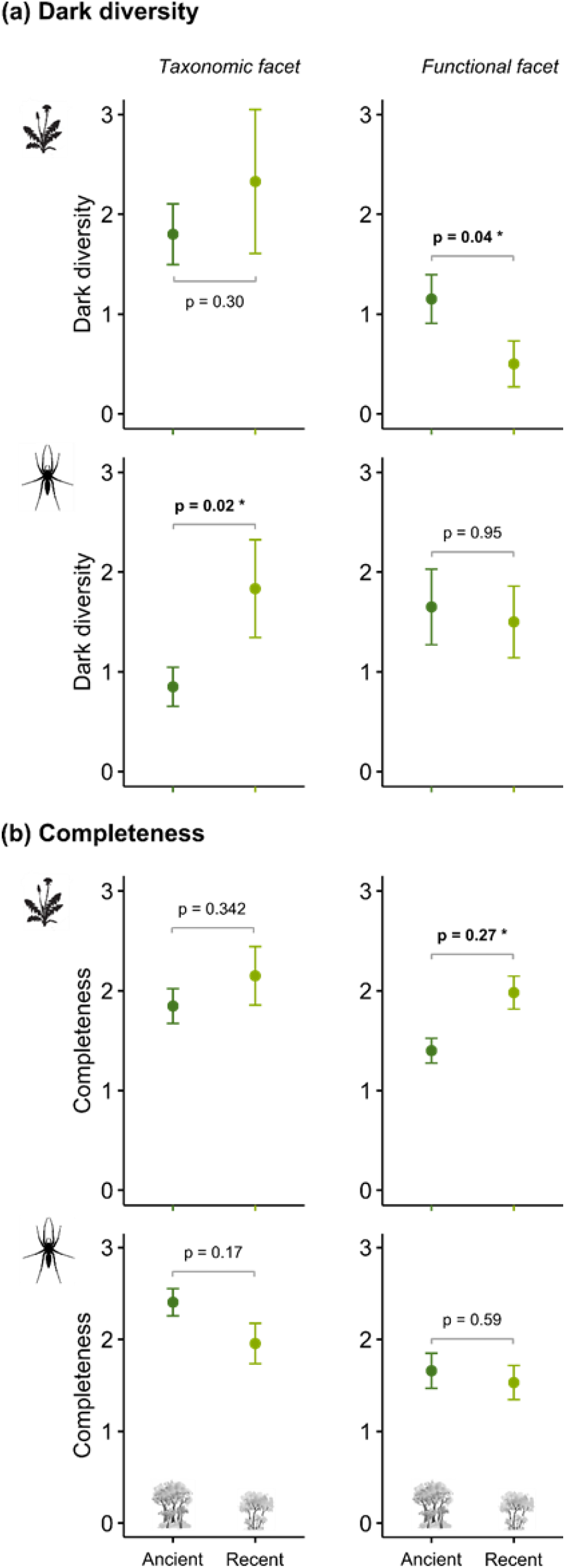
Comparisons of dark diversity (a) and completeness (b) between ancient and recent forests, for plant and spider communities, and for the taxonomic and the functional facets (n=32).

## Discussion

Our application of the dark diversity framework into a case study of passive rewilding revealed several novel ecological insights concerning the mechanisms involved in the re-assembly of natural communities during land-use changes. Moreover, the direct quantification of functional dark diversity brought new light on the potential abilities of recent woodlands to spontaneously recovering native forest biodiversity. We thus demonstrated that, surprisingly, recent forests were in fact quite complete from an overall, ecological perspective. Despite lacking specialist, plant and spider species, recent woodlands already harboured functionally rich communities.

### Taxonomic and functional dark diversity: two distinct but complementary facets of potential biodiversity

Our results obtained from functional dark diversity clearly differed from those obtained from taxonomic dark diversity, both in terms of quality (*i.e.* composition of dark diversity) and quantity (*i.e.* amount of dark diversity). For plants and spiders, the taxonomical approach integrated only one quarter of all species into dark diversity, whereas about half of all functional groups were integrated at least once into functional dark diversity. Thus, our results suggest that an exclusively taxonomic approach tends in fact to overestimate the ecological integrity of communities, by missing out the fact that some niches are actually vacant in several communities. By focusing on the co-occurrence of functional groups rather than co-occurrence of species, and considering that *any* species of a given lacking group could be recruited, our approach enabled to identify vacant niches even when the species of the group concerned were not integrated into taxonomic dark diversity. For example, no species of short-lived herbs (FG #4) or medium-size generalist hunting spiders (FE #4) was ever integrated into taxonomic dark diversity, while these groups were often integrated into functional dark diversity (Figure 3). We thus note that our approach increased the probability of detecting the absence of functional features hosted by several rare species, whereas neither taxonomic dark diversity nor a functional interpretation of it could detect them (as we assumed, see Figure 1d).

On the other hand, functional dark diversity as we calculated it might, too, overestimate the ecological integrity of natural communities, because it considers communities represented by a single species per group as complete (see Figure 1b). Thus, species may be lacking but their respective groups may not: for example, shrubs and ferns were never included into dark diversity, whereas some of their species were (Figure 3). We could hence summarise our approach in simple words: functional dark diversity is not interested in species. This can be a major advantage: for habitat conservation and restoration, it is often crucial to investigate ecosystem functioning and related services before assessing their richness or originality in species (Cadotte, Carscadden, & Mirotchnick, 2011). However, this could be a drawback in other cases: conservation and restoration policies can also target species for their intrinsic patrimonial value (e.g. existence values), hence requiring consideration of species. Rare species may also play a key role in ecosystem functioning by ensuring singular functions or enhancing functional redundancy (Chapman, Tunnicliffe, & Bates, 2018; Leitão et al., 2016). We hence suggest that further methods need to be developed to measure the potential regeneration of natural habitats considering rare species with rare functional features. Overall, we think that taxonomic dark diversity and functional dark diversity illustrate different facets of communities, and that they can be very complementary metrics which, taken together, provide reliable information for ecological diagnostic and for conservation and restoration policies.

### Dark diversity brings to light recruitment limitations during the forest recovering process but with little impact on forest functional integrity

Our results confirmed that recent woodlands, even after decades of forest re-establishing, do not fully recover communities like those of ancient forests in terms of species identity, whether for plants (Bergès & Dupouey, 2020; Hermy & Verheyen, 2007) or spiders (Morel et al., 2019). Dark diversity showed that recent forests mainly lacked some generalist species they could recruit (e.g. phanerophytes or ruderal-nitrophilous plants such as *Crataegus monogyna* and *Ajuga reptans*, and several ubiquitous hunter spiders such as *Agroeca brunnea* and *Pardosa saltans*). Moreover, we observed that recent forests also lacked specialist forest species, but they might not be able to be recruited, since they were not identified in dark diversity. These specialists, which are mainly slow- colonisers associated with specific, restricted ecological conditions (e.g. oligotrophic and acidophil soils for plants, Hermy & Verheyen, 2007, and complex litters associated to dead-wood materials for spiders, Morel et al., 2019) were almost exclusively associated, when they were absent, to ancient forests. Therefore, all these compositional differences in the dark diversity suggest that a recruitment limitation due to past land-uses was, here, the main driver of the reshaping of communities, rather than a dispersal limitation.

Beyond these changes in species identities, dark diversity also showed that recent woodlands harboured diverse communities which were quite complete from a functional perspective, especially regarding plants. Recent woodlands mainly lacked small springs herbs whereas ancient forests also lacked shrubs, vernal geophytes and various herbs. Three complementary hypotheses could explain this result. First, recent woodlands might temporary harbour “relictual species” (and their functional features) inherited from preceding successional stages (e.g. shrubs species), which might be in extinction debt and could disappear with time (Bagaria, Helm, Rodà, & Pino, 2015). Second, past land uses may have reduced nutrient limitation through soil fertilisation, particularly on the acid soils of our study region (Graae, 2000; Koerner et al., 1997), leading to recruitment of more diverse functional features in recent woodlands (Morel et al., 2019b). Last, past management of forests might also play a role: since several centuries, the management of ancient forests has shifted from coppicing to high-forest system, which has tended to disadvantage shade-loving, understorey woody and herb species (Kirby & Watkins, 2015). On the contrary, recent woodlands conserve a denser coppice, thus leading to a lower amount of light reaching their understorey (illustrated by the Ellenberg values, Table S1), which could enrich the herbaceous cover in both species and functional plant features. According to these last two hypotheses, compositional differences between both forest types should be maintained with time.

Overall, we believe that our results may challenge and improve our perception of the conservation value of both recent and ancient forests: recent woodlands do lack typical ancient forest species, but they can also recover functionally rich and ecologically complete communities. Even if forestry is not incompatible with biodiversity, we think that an increase in wildwood areas could benefit to conservation of forest ecosystems.

### Perspectives and limitations

We argue for the development of the framework of functional dark diversity for both researchers and practicioners, notably in the study of biodiversity responses to land-use changes. First, we acknowledge some limitations of our results: our recent study forests might be in somewhat good conditions compared to other ones elsewhere in the study region, because they have not undergone a particularly excessive anthropogenic pressure during their regeneration. We also note that we studied dark diversity on a relatively small dataset (*i.e.* several forests of Brittany), but we think it was sufficiently robust to analyse the different facets of dark diversity and test their dissemblances. In addition, the fact that both compositional and diversity patterns are congruent between the two distinct taxa studied (especially in term of functional integrity within recent woodlands), tends to confirm the robustness of our results. Overall, we think that our method assessing functional dark diversity, with its simplicity, can easily be applied to many other issues of conservation and restoration. The combined use of functional and taxonomic dark diversity can deal with the assessment of the ecological integrity of natural communities, both from a functional perspective (including resistance and resilience capacities of ecosystems) and from a taxonomic one (e.g. recruitment of species with particular interest). Since the method is entirely based on the co-occurrence of functional groups, the choice of these functional groups is a central concern. We ensured that functional groups satisfied two conditions: functional redundancy had to be higher within groups than between groups, and coexistence within groups had to be neutralist. We also ensured that the selected functional groups corresponded to a precise ecological compartment, that is, a subset of species that was noticeable in the field. In this way, we think that the functional group approach for dark diversity may be used, and does have a biological sense. Further methods could be developed in the future, using a continuous approach for traits along the whole calculation process, or focusing on the specialisation degree, the evolutionary distinctiveness or the functional originality of species present in the dark diversity.

## Acknowledgements

We are very grateful to the “Région Bretagne”, the “Conseil départemental des Côtes d’Armor”, the “Conseil départemental d’Ille-et-Vilaine” and the “Communauté de communes de Plouha-Lanvollon” which supported financially this work. We also are very grateful to Benoit Dujol for fruitful early discussion about dark diversity.

## Supplementary Information

**Table S1.** Mean values of environmental parameters in recent woodlands and ancient forests (mean ± standard deviation). Habitat structure was assessed from canopy cover (visual vertical estimation above each plot) and basal area (measured with a chain-relascope). We infer abiotic conditions from Ellenberg Indicator Values (EIV) for moisture, reaction, nitrogen and light, using flora data adapted to the Western Europe flora (Hill 1999). EIV were weighted by the vegetation cover (in %) to account for species abundances. We determined differences using Student tests.

|  | Recent woodlands | Ancient forests | *p*-value |
| --- | --- | --- | --- |
| *n* plots | 12 | 20 |  |
| **Habitat structure** |  |  |  |
| Canopy cover (%) | 74.6 ± 3.1 | 77.3 ± 3.1 | 0.549 |
| Basal area (m<sup>2</sup> ha<sup>-1</sup>) | 27.6 ± 1.9 | 31.1 ± 1.9 | 0.217 |
| **Ellenberg Indicator Value (EIV)** |  |  |  |
| Moisture (F) | 5.65 ± 0.04 | 5.62 ± 0.04 | 0.54 |
| Reaction (R) | 5.93 ± 0.08 | 4.62 ± 0.1 | < 0.001 |
| Nitrogen (N) | 5.72 ± 0.06 | 4.52 ± 0.1 | < 0.001 |
| Light (L) | 5.12 ± 0.08 | 5.42 ± 0.06 | 0.008 |

Hill, M.O., 1999. Ellenberg's Indicator Values for British Plants. DETR, Rotherham.

**Table S2.** Description of functional traits used to characterise plant species, from the LEDA database. Trait types: 1=quantitative, 2=qualitative, 3=ordinal.

|  | Functional traits | Type | Description and units |
| --- | --- | --- | --- |
| 1 | Dispersal syndrome<br>99 (100%) | 2 | Anemochory; autochory; barochory; hydrochory; endozoochory; epizoochory and myrmecochory |
| 2 | Leaf dry matter content (LDMC)<br>99 (95%) | 1 | mg.g<sup>-1</sup> |
| 3 | Plant height (Height)<br>99 (100%) | 1 | Maximum vegetative height (m) |
| 4 | Plant life-form<br>99 (100%) | 2 | Geophyte; hemicryptophyte, liana-phanerophyte and phanerophyte |
| 5 | Plant life-span<br>99 (100%) | 1 | Perennial and short-lived |
| 6 | Seed dry mass<br>89 /99 (89%) | 1 | mg |
| 7 | Pollination<br>99 (100%) | 2 | Anemogamy; autogamy; entomogamy; hydrogamy |
| 8 | Specific leaf area (SLA)<br>99 (100%) | 1 | mm<sup>2</sup>. g<sup>-1</sup> |
| 9 | Flowering phenology<br>99 (100%) | 3 | Start of flowering: months (1-12) |

**Table S3.** Description of functional traits used to characterize spider species, from the literature.

| Functional traits | Description and units | Reference |
| --- | --- | --- |
| Body size | Categorical (3)<br>Small (<9 mm); Medium (9-15mm);<br>Large (> 15mm) | Roberts (1995) |
| Guilds | Categorical (7)<br>Ambush hunters, Ground hunters,<br>Others hunters, Sheet web, Space<br>web, Orb web, Wandering web | Cardoso et al., (2011) |
| Phenology | Categorical (4)<br>Spring, Summer, Autumn-Winter,<br>Eurychron | Harvey et al., (2002) |
| Circadian activity | Categorical (3)<br>Diurnal, Nocturnal, Both | Cardoso et al., (2011) |

Buchar, J. & Ruzicka, V. (2002) *Catalogue of Spiders of the Czech Republic*.
Cardoso, P., Pekár, S., Jocqué, R. & Coddington, J.A. (2011) Global Patterns of Guild Composition and
Functional Diversity of Spiders (ed M Somers). *PLoS ONE*, 6, e21710.
Harvey, P.R., Nellist, D.R. & Telfer, M.G. (2002) *Provisional Atlas of British Spiders (Arachnida,*
*Araneae)*, Biological Records Center. Huntington, UK.
Roberts, M.J. (1995) *Spiders of Britain & Northern Europe*. Harper Collins, London; New York.

**Table S4.** Description of functional entities (FE) identified for spiders.

| # FE | Functional entities | Body | Phenology | Circadian activity | Hunting | n species |
| --- | --- | --- | --- | --- | --- | --- |
| 1 | Bla_Peu_Dbo_Hgr | Large | Eurychron | Both | Ground_hunters | 3 |
| 2 | Bsm_Psu_Dbo_Hsh | Small | Summer | Both | Sheetweb | 1 |
| 3 | Bla_Psu_Ddi_Hgr | Large | Summer | Diurnal | Ground_hunters | 6 |
| 4 | Bme_Peu_Dbo_Hgr | Medium | Eurychron | Both | Ground_hunters | 2 |
| 5 | Bla_Peu_Dbo_Ham | Large | Eurychron | Both | Ambush_hunters | 1 |
| 6 | Bsm_Peu_Dbo_Hsh | Small | Eurychron | Both | Sheetweb | 10 |
| 7 | Bsm_Pau_Dbo_Hsh | Small | Autumn-Winter | Both | Sheetweb | 4 |
| 8 | Bme_Peu_Dbo_Hsh | Medium | Eurychron | Both | Sheetweb | 6 |
| 9 | Bsm_Psu_Dbo_Hwa | Small | Summer | Both | Wandering_web | 4 |
| 10 | Bla_Psu_Dno_Hot | Large | Summer | Nocturnal | Otherhunters | 1 |
| 11 | Bla_Peu_Dno_Hot | Large | Eurychron | Nocturnal | Otherhunters | 1 |
| 12 | Bla_Psp_Dno_Hsh | Large | Spring | Nocturnal | Sheetweb | 1 |
| 13 | Bsm_Peu_Dbo_Hwa | Small | Eurychron | Both | Wandering_web | 15 |
| 14 | Bla_Psp_Dbo_Hgr | Large | Spring | Both | Ground_hunters | 1 |
| 15 | Bla_Peu_Dno_Hgr | Large | Eurychron | Nocturnal | Ground_hunters | 1 |
| 16 | Bme_Peu_Dbo_Hsp | Medium | Eurychron | Both | Spaceweb | 2 |
| 17 | Bla_Pau_Dbo_Hsh | Large | Autumn-Winter | Both | Sheetweb | 1 |
| 18 | Bme_Peu_Dbo_Hwa | Medium | Eurychron | Both | Wandering_web | 5 |
| 19 | Bme_Psu_Dbo_Hwa | Medium | Summer | Both | Wandering_web | 1 |
| 20 | Bla_Psu_Dbo_Hgr | Large | Summer | Both | Ground_hunters | 3 |
| 21 | Bla_Psp_Ddi_Hgr | Large | Spring | Diurnal | Ground_hunters | 1 |
| 22 | Bla_Peu_Dbo_Hsh | Large | Eurychron | Both | Sheetweb | 1 |
| 23 | Bme_Psp_Dbo_Hsh | Medium | Spring | Both | Sheetweb | 1 |
| 24 | Bla_Psu_Dbo_Hor | Large | Summer | Both | Orbweb | 1 |
| 25 | Bsm_Peu_Ddi_Hot | Small | Eurychron | Diurnal | Otherhunters | 1 |
| 26 | Bme_Peu_Dbo_Ham | Medium | Eurychron | Both | Ambush_hunters | 2 |
| 27 | Bla_Peu_Dbo_Hor | Large | Eurychron | Both | Orbweb | 1 |
| 28 | Bme_Peu_Dbo_Hor | Medium | Eurychron | Both | Orbweb | 1 |
| 29 | Bla_Peu_Ddi_Hgr | Large | Eurychron | Diurnal | Ground_hunters | 3 |
| 30 | Bsm_Psu_Dbo_Hgr | Small | Summer | Both | Ground_hunters | 1 |
| 31 | Bsm_Psp_Dbo_Hwa | Small | Spring | Both | Wandering_web | 2 |
| 32 | Bsm_Pau_Dbo_Hwa | Small | Autumn-Winter | Both | Wandering_web | 1 |
| 33 | Bsm_Psu_Dbo_Hsp | Small | Summer | Both | Spaceweb | 1 |
| 34 | Bme_Psp_Dbo_Hwa | Medium | Spring | Both | Wandering_web | 1 |
| 35 | Bla_Psu_Dbo_Ham | Large | Summer | Both | Ambush_hunters | 2 |

**Table S5.**
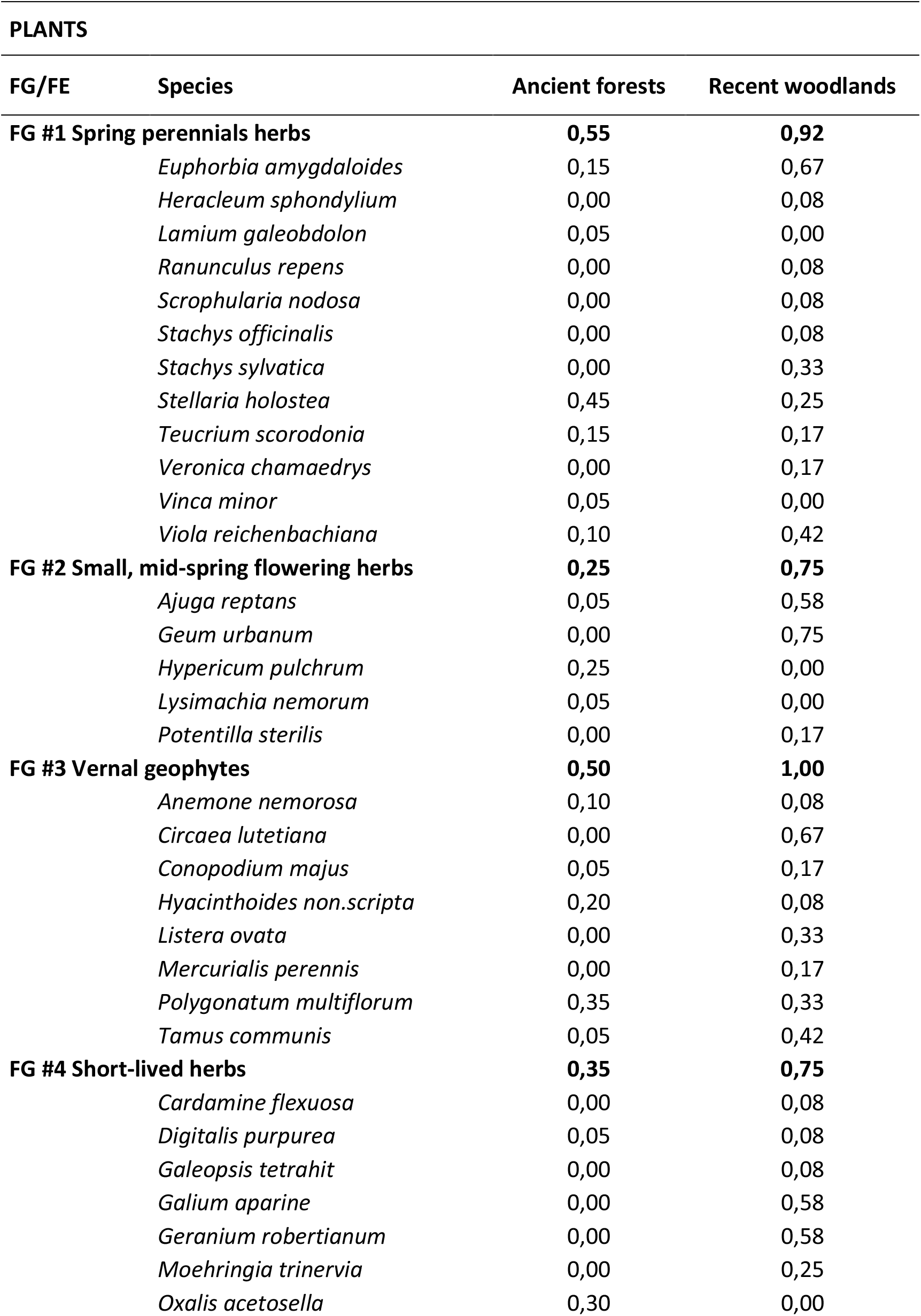
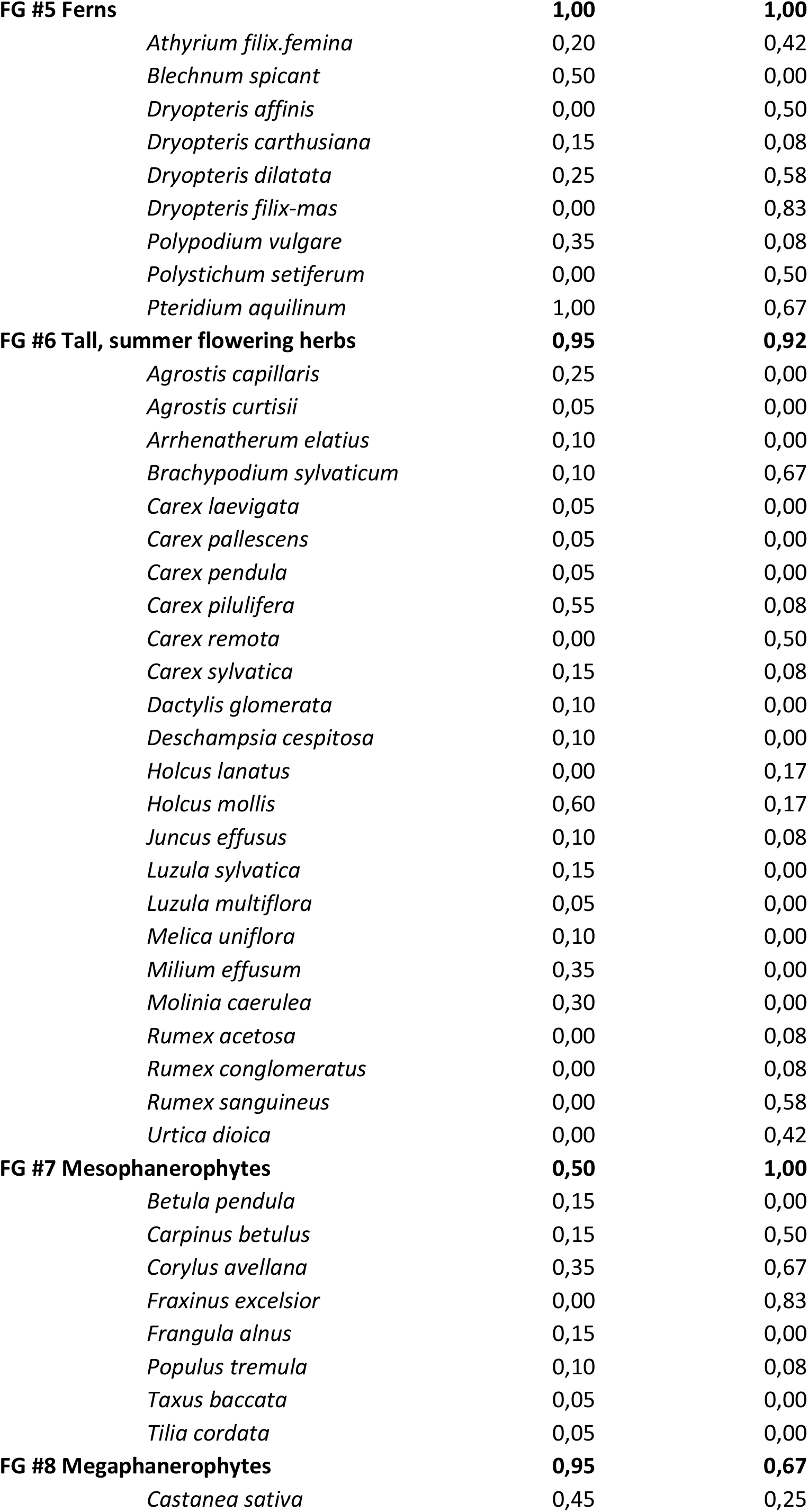
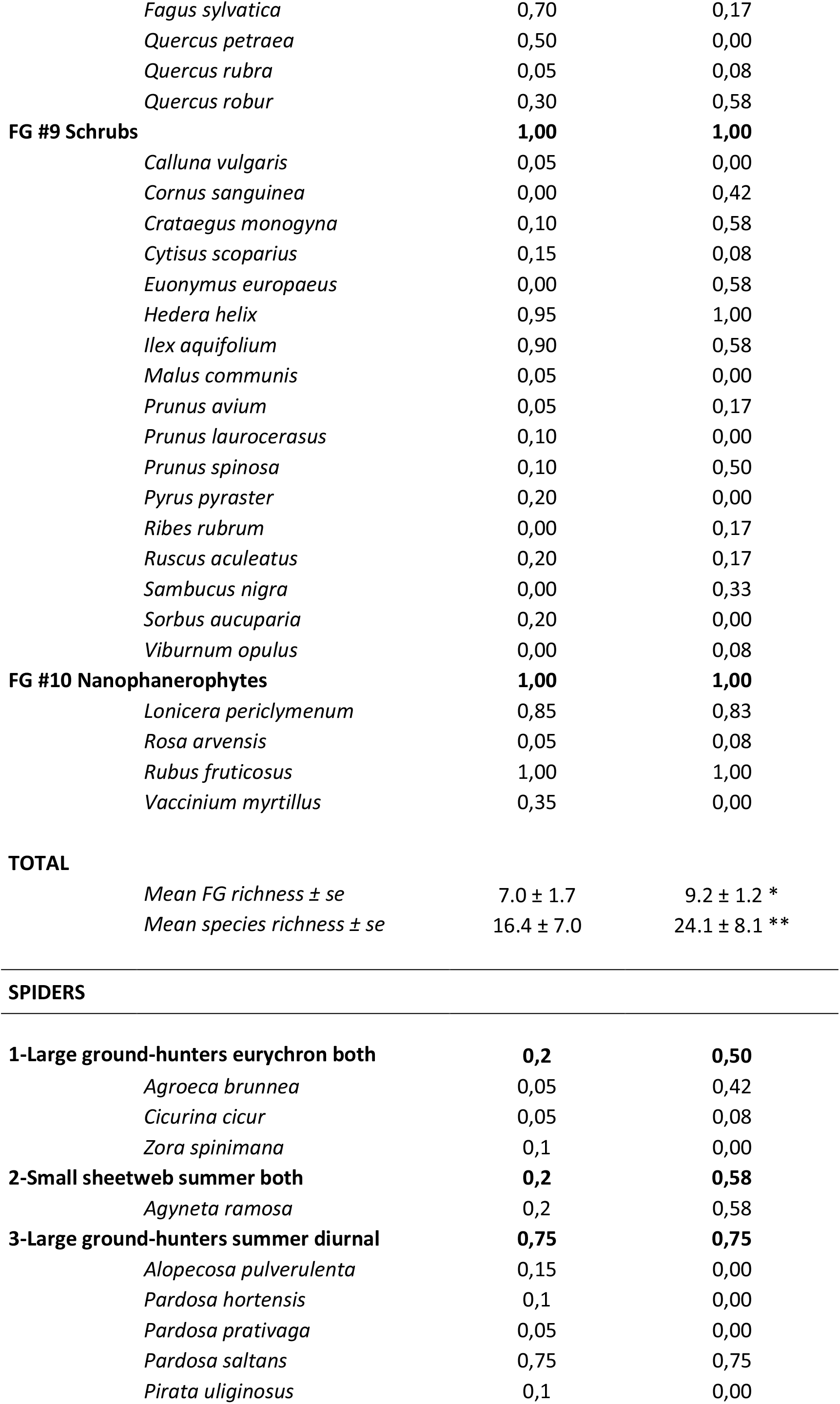
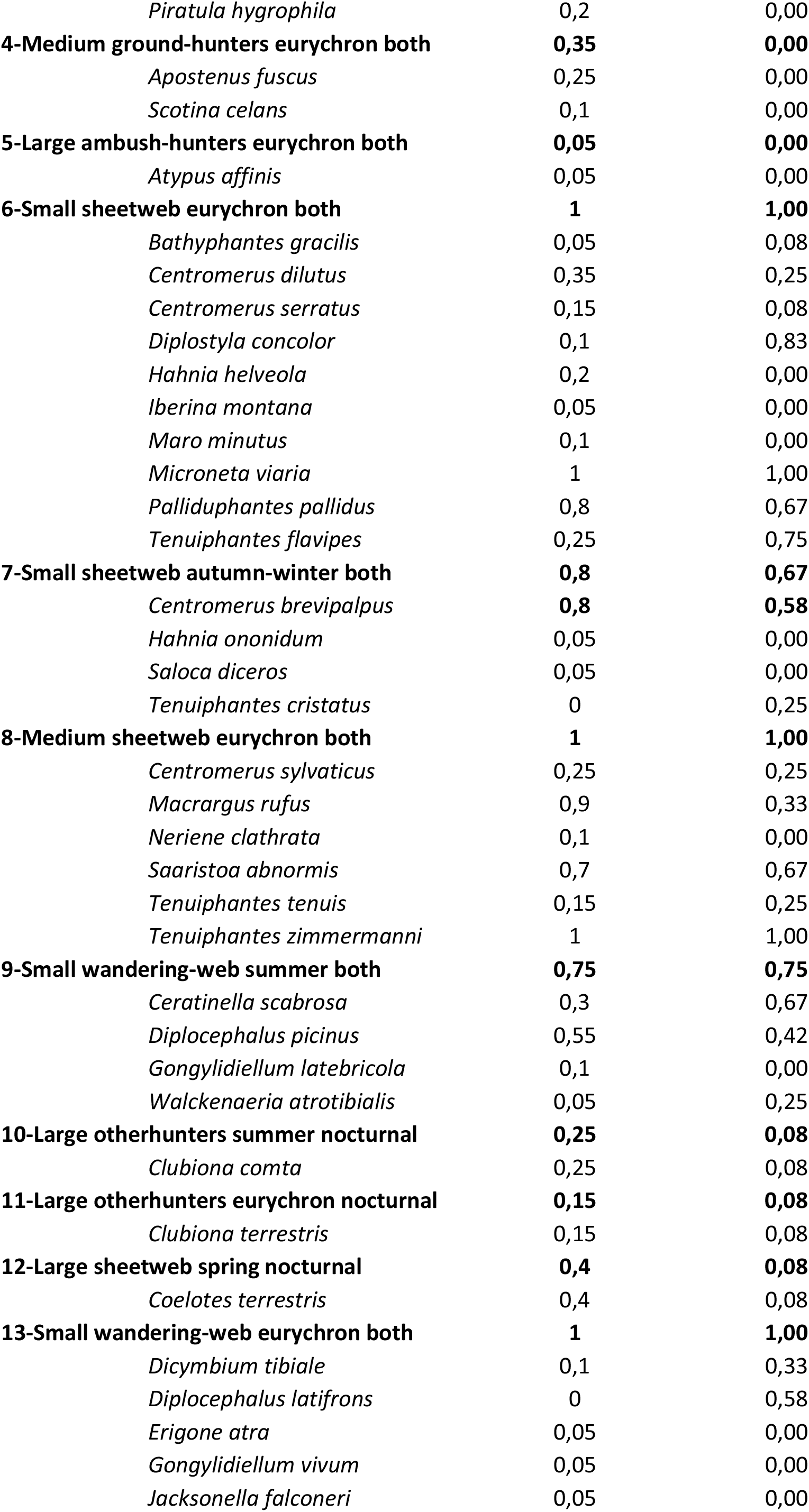
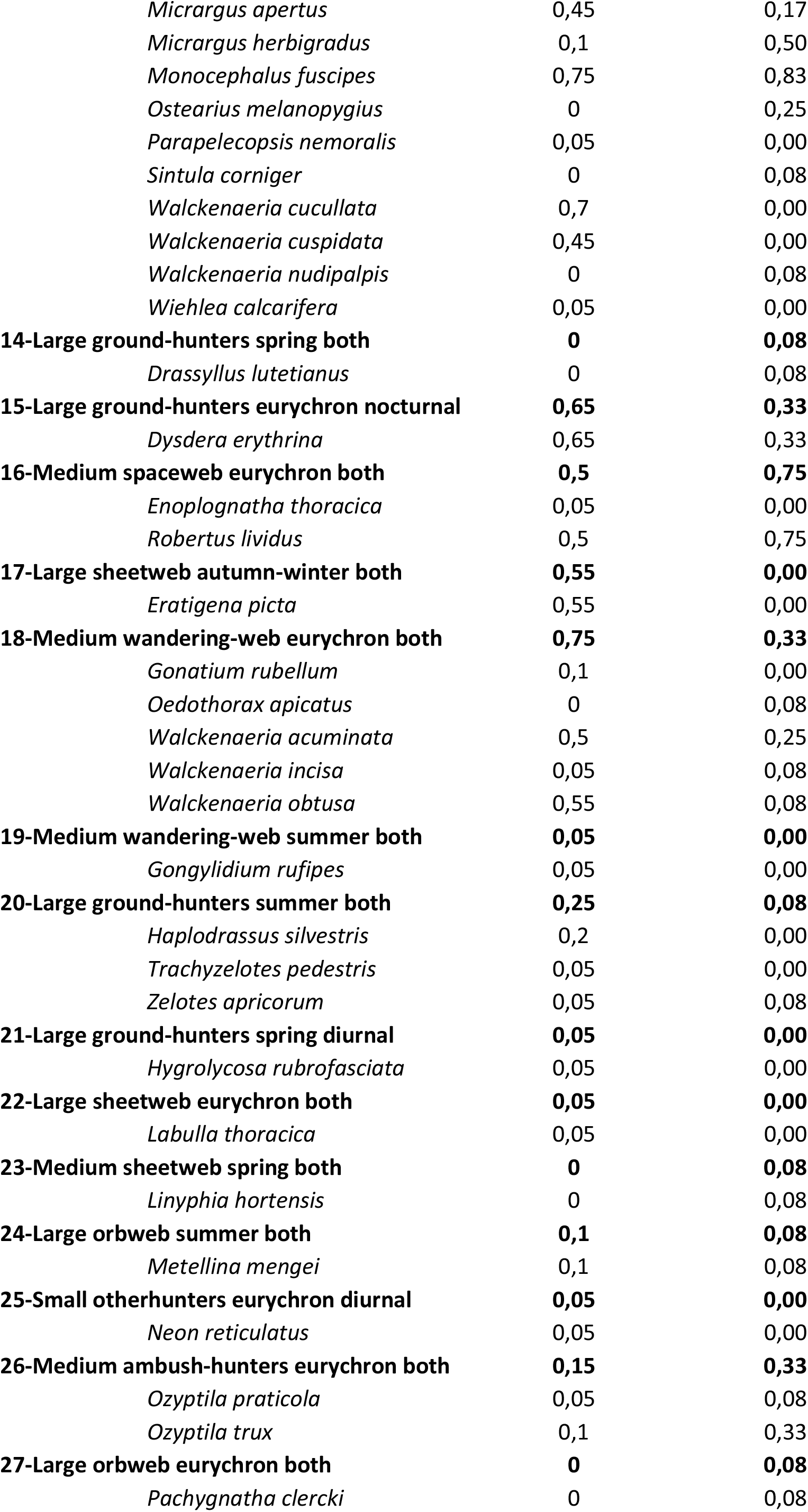
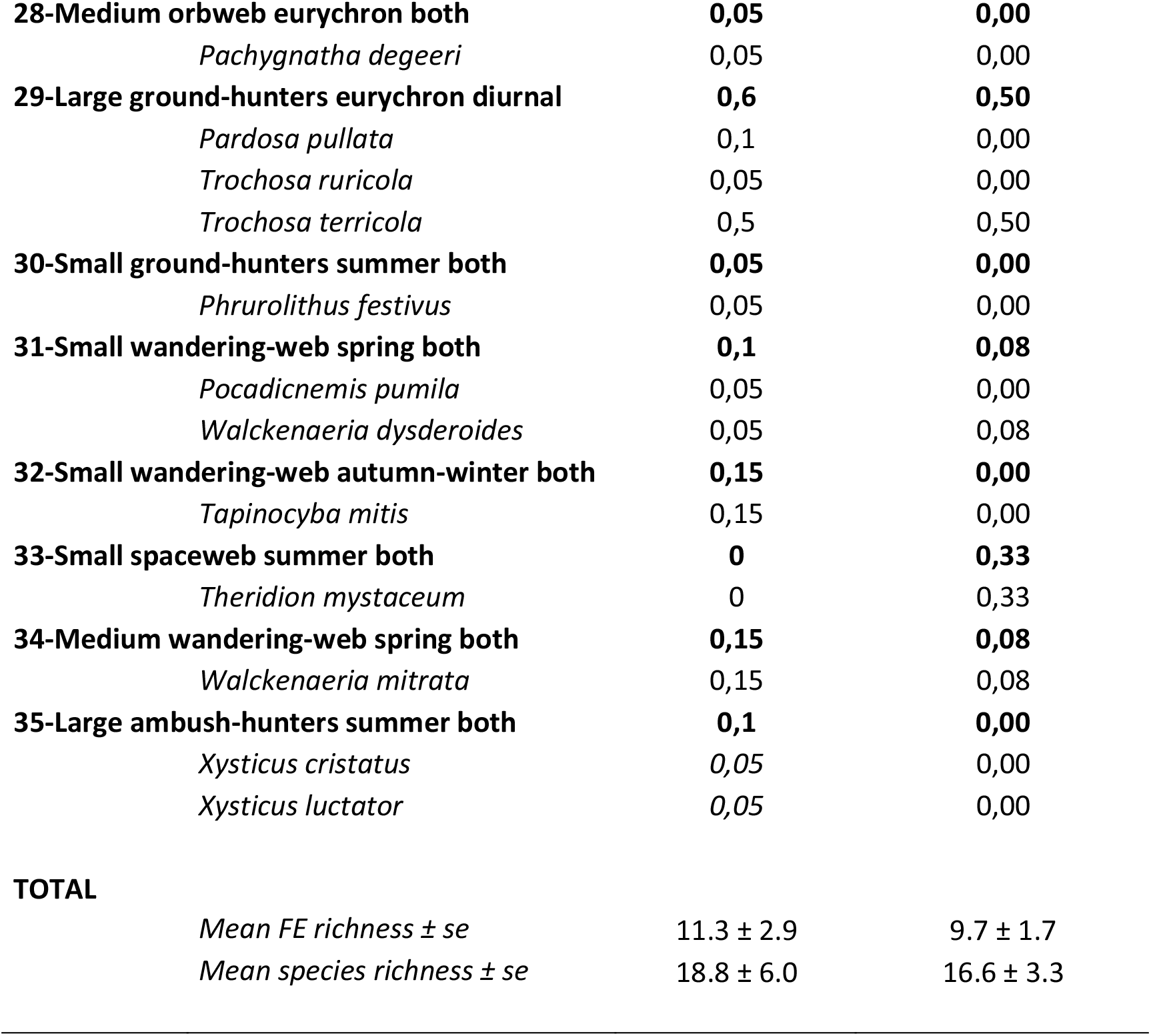
Occurrence frequency (%) in ancient and recent forests of observed diversity of (i) functional groups of plants (in bold) and plant species and (ii) functional entities of spiders (in bold) and spider species. The average observed diversity per plot is also given, that is, the average richness per plot in terms of species and functional groups and entities. See Table S4 and Figure S1 for description of functional groups end entities.

**Figure S1.**
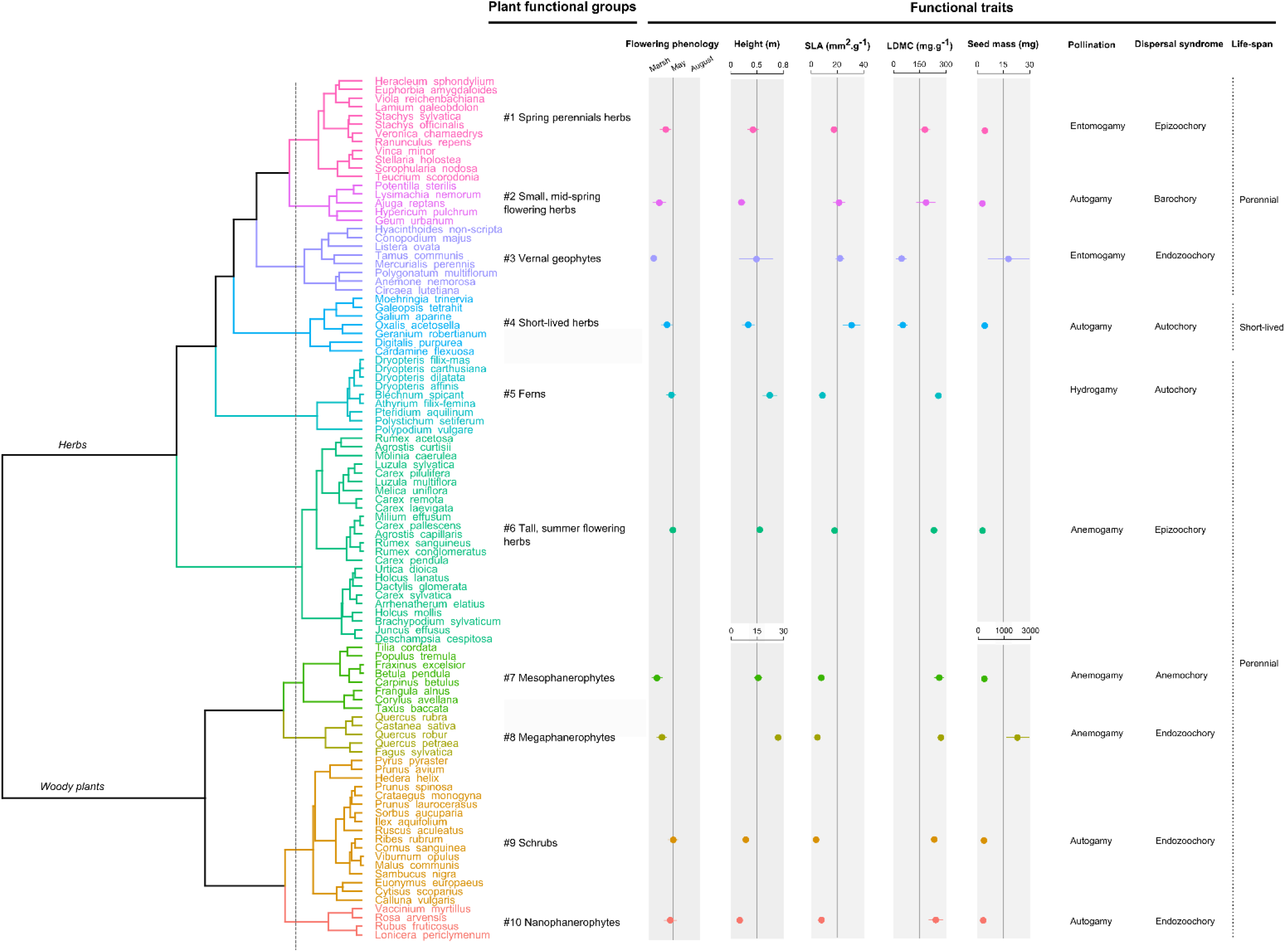
Functional groups of plant species identified with the dendrogram method (see Materials and Methods for more details). For each functional group, the mean of continuous traits and the dominant modality of categorical traits are given.

**Figure S2.**
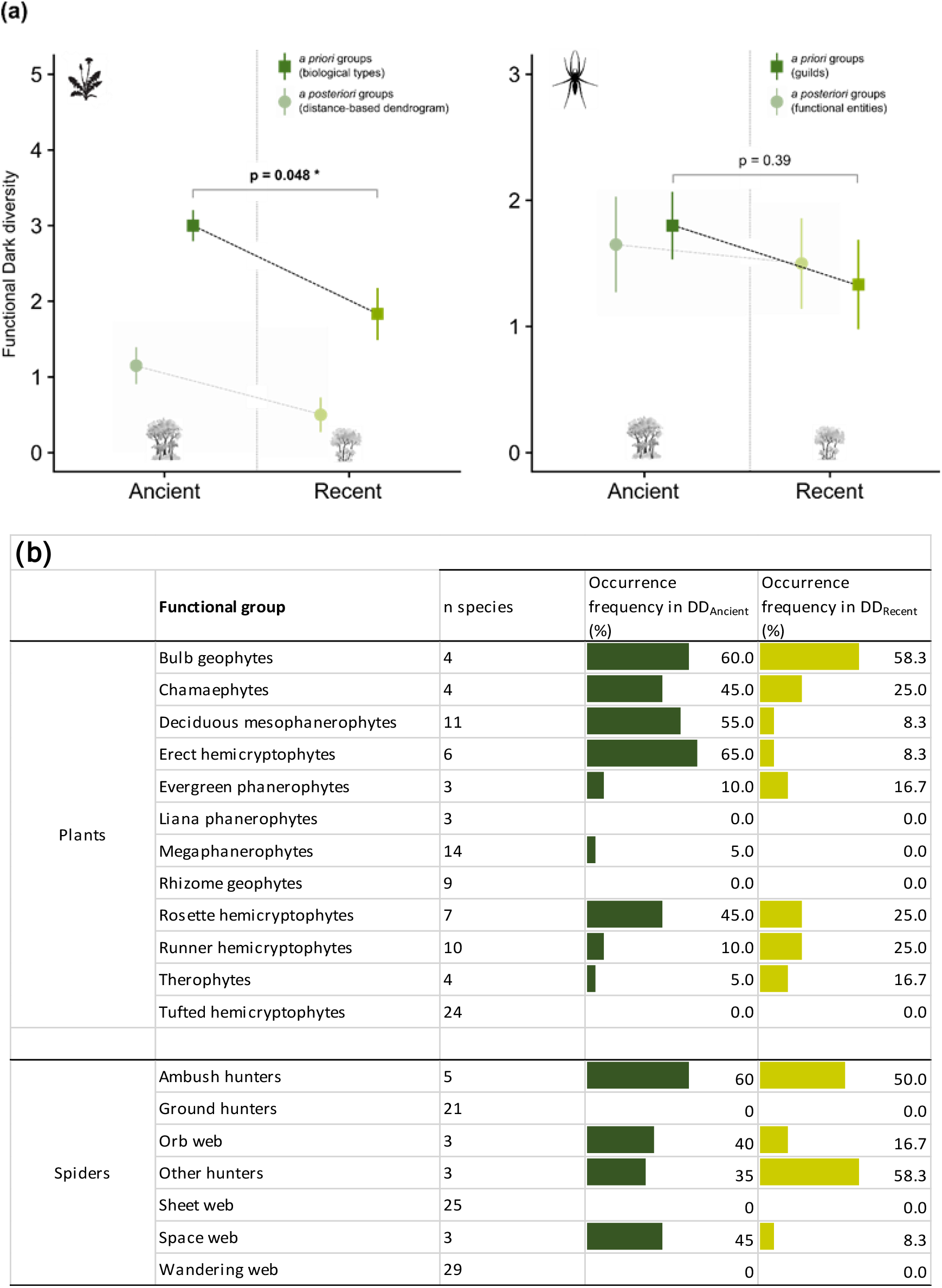
Comparisons between ancient and recent forests of (a) functional dark diversity and (b) its composition, based on functional groups defined *a priori* (Julve’s groups for plants and Cardoso’s guilds for spiders). Results are very similar to those obtained with groups defined *a posteriori* (Figure 4 and in light grey here): functional dark diversity is higher in ancient forests for plant communities (and with a different composition), and functional dark diversity is equivalent between both forest types for spiders (with also a different composition).

## References

Bagaria, G., Helm, A., Rodà, F., & Pino, J. (2015). Assessing coexisting plant extinction debt and colonization credit in a grassland–forest change gradient. Oecologia, 179(3), 823–834.

Beals, A. (1984). Bray-Curtis ordination: an effective strategy for analysis of multivariate ecological data. Advances in Ecological Research, 14, 1–55.

Bergès, L., & Dupouey, J. (2020). Historical ecology and ancient forests: Progress, conservation issues and scientific prospects, with some examples from the French case. Journal of Vegetation Science, jvs.12846.

Cadotte, M. W., Carscadden, K., & Mirotchnick, N. (2011). Beyond species: functional diversity and the maintenance of ecological processes and services: Functional diversity in ecology and conservation. Journal of Applied Ecology, 48(5), 1079–1087.

Cardoso, P., Pekár, S., Jocqué, R., & Coddington, J. A. (2011). Global Patterns of Guild Composition and Functional Diversity of Spiders. PLoS ONE, 6(6), e21710. doi: 10.1371/journal.pone.0021710

Cateau, E., Larrieu, L., Vallauri, D., Savoie, J.-M., Touroult, J., & Brustel, H. (2015). Ancienneté et maturité: deux qualités complémentaires d’un écosystème forestier. Comptes Rendus Biologies, 338(1), 58–73.

Chapman, A. S. A., Tunnicliffe, V., & Bates, A. E. (2018). Both rare and common species make unique contributions to functional diversity in an ecosystem unaffected by human activities. Diversity and Distributions, 24(5), 568–578.

Chase, J. M., & Leibold, M. A. (2003). Ecological niches: linking classical and contemporary approaches. Chicago: University of Chicago Press.

Flinn, K. M., & Vellend, M. (2005). Recovery of forest plant communities in post-agricultural landscapes. Frontiers in Ecology and the Environment, 3(5), 243–250.

Graae, B. J. (2000). The effect of landscape fragmentation and forest continuity on forest floor species in two regions of Denmark. Journal of Vegetation Science, 11(6), 881–892.

Helm, A., Zobel, M., Moles, A. T., Szava-Kovats, R., & Pärtel, M. (2015). Characteristic and derived diversity: implementing the species pool concept to quantify conservation condition of habitats. Diversity and Distributions, 21(6), 711–721.

Hermant, M., Hennion, F., Bartish, I., Yguel, B., & Prinzing, A. (2012). Disparate relatives: Life histories vary more in genera occupying intermediate environments. Perspectives in Plant Ecology, Evolution and Systematics, 14, 283–301.

Hermy, M., & Verheyen, K. (Eds.). (2007). Legacies of the past in the present-day forest biodiversity: a review of past land-use effects on forest plant species composition and diversity. In Sustainability and Diversity of Forest Ecosystems: an Interdisciplinary Approach (pp. 361–371). Tokyo; New York: Springer.

Hermy, Martin, Honnay, O., Firbank, L., Grashof-Bokdam, C., & Lawesson, J. E. (1999). An ecological comparison between ancient and other forest plant species of Europe, and the implications for forest conservation. Biological Conservation, 91(1), 9–22.

Hofmeister, J., Hošek, J., Brabec, M., Hermy, M., Dvořák, D., Fellner, R., … Kadlec, T. (2019) Shared affinity of various forest-dwelling taxa point to the continuity of temperate forests. Ecological Indicators, 101, 904–912.

Jackson, S. T., & Sax, D. F. (2010). Balancing biodiversity in a changing environment: extinction debt, immigration credit and species turnover. Trends in Ecology & Evolution, 25(3), 153–160.

Julve, P. (1998). Baseflor: index botanique, écologique et chorologique de la flore de France. Institut Catholique de Lille.

Kimberley, A., Blackburn, G. A., Whyatt, J. D., Kirby, K., & Smart, S. M. (2013). Identifying the trait syndromes of conservation indicator species: how distinct are British ancient woodland indicator plants from other woodland species? Applied Vegetation Science, 16(4), 667–675.

Kirby, K., Watkins, C., 2015. Europe’s Changing Woods and Forests: From Wildwood to Managed Landscapes. CABI, 363p.

Kleyer, M., Bekker, R. M., Knevel, I. C., Bakker, J. P., Thompson, K., Sonnenschein, M., … Peco, B. (2008). The LEDA Traitbase: a database of life-history traits of the Northwest European flora. Journal of Ecology, 96(6), 1266–1274.

Koerner, W., Dupouey, J. L., Dambrine, E., & Benoit, M. (1997). Influence of Past Land Use on the Vegetation and Soils of Present Day Forest in the Vosges Mountains, France. The Journal of Ecology, 85(3), 351.

Lavorel, S., & Garnier, E. (2002). Predicting changes in community composition and ecosystem functioning from plant traits: revisiting the Holy Grail. Functional Ecology, 16(5), 545–556.

Leitão, R. P., Zuanon, J., Villéger, S., Williams, S. E., Baraloto, C., Fortunel, C., … Mouillot, D. (2016). Rare species contribute disproportionately to the functional structure of species assemblages. Proceedings of the Royal Society B: Biological Sciences, 283(1828), 20160084.

Lewis, R. J., de Bello, F., Bennett, J. A., Fibich, P., Finerty, G. E., Götzenberger, L., … Pärtel, M. (2017). Applying the dark diversity concept to nature conservation: Dark Diversity and Nature Conservation. Conservation Biology, 31(1), 40–47.

Lewis, R. J., Szava-Kovats, R., & Pärtel, M. (2016). Estimating dark diversity and species pools: an empirical assessment of two methods. Methods in Ecology and Evolution, 7(1), 104–113.

Mcgill, B., Enquist, B., Weiher, E., & Westoby, M. (2006). Rebuilding community ecology from functional traits. Trends in Ecology & Evolution, 21(4), 178–185.

Moeslund, J. E., Brunbjerg, A. K., Clausen, K. K., Dalby, L., Fløjgaard, C., Juel, A., & Lenoir, J. (2017). Using dark diversity and plant characteristics to guide conservation and restoration. Journal of Applied Ecology, 54(6), 1730–1741.

Morel, L., Dujol, B., Courtial, C., Vasseur, M., Leroy, B., & Ysnel, F. (2019a). Spontaneous recovery of functional diversity and rarity of ground-living spiders shed light on the conservation importance of recent woodlands. Biodiversity and Conservation, 28(3), 687–709.

Morel, L., Barbe, L., Jung, V., Clément, B., Schnitzler, A. & Ysnel, F. (2019b). Passive rewilding may (aslo) restore phylogentically rich and functionally resilient forest plant communities. Ecological Applications, 30(1), e02007.

Mouillot, D., Villeger, S., Parravicini, V., Kulbicki, M., Arias-Gonzalez, J. E., Bender, M., … Bellwood, D. R. (2014). Functional over-redundancy and high functional vulnerability in global fish faunas on tropical reefs. Proceedings of the National Academy of Sciences, 111(38), 13757–13762.

Murtagh, F., & Legendre, P. (2014). Ward’s Hierarchical Agglomerative Clustering Method: Which Algorithms Implement Ward’s Criterion? Journal of Classification, 31(3), 274–295.

Newbold, T., Hudson, L. N., Hill, S. L. L., Contu, S., Lysenko, I., Senior, R. A., … Purvis, A. (2015). Global effects of land use on local terrestrial biodiversity. Nature, 520(7545), 45–50.

Oxbrough, A. G., Gittings, T., O’Halloran, J., Giller, P. S., & Smith, G. F. (2005). Structural indicators of spider communities across the forest plantation cycle. Forest Ecology and Management, 212(1–3), 171–183.

Pärtel, M. (2014). Community ecology of absent species: hidden and dark diversity. Journal of Vegetation Science, 25(5), 1154–1159.

Pärtel, M., Szava-Kovats, R., & Zobel, M. (2011). Dark diversity: shedding light on absent species. Trends in Ecology & Evolution, 26(3), 124–128.

Pärtel, M., Szava-Kovats, R., & Zobel, M. (2013). Community Completeness: Linking Local and Dark Diversity within the Species Pool Concept. Folia Geobotanica, 48(3), 307–317.

Pérez-Harguindeguy, N., Díaz, S., Garnier, E., Lavorel, S., Poorter, H., Jaureguiberry, P., … Cornelissen, J. H. C. (2013). New handbook for standardised measurement of plant functional traits worldwide. Australian Journal of Botany, 61(3), 167.

Prinzing, A., Reiffers, R., Braakhekke, W. G., Hennekens, S. M., Tackenberg, O., Ozinga, W. A., … van Groenendael, J. M. (2008). Less lineages more trait variation: phylogenetically clustered plant communities are functionally more diverse. Ecology Letters, 11(8), 809–819.

Verheyen, K., Honnay, O., Motzkin, G., Hermy, M., & Foster, D. R. (2003). Response of forest plant species to land-use change: a life-history trait-based approach. Journal of Ecology, 91(4), 563–577.

Violle, C., Navas, M.-L., Vile, D., Kazakou, E., Fortunel, C., Hummel, I., & Garnier, E. (2007). Let the concept of trait be functional! Oikos, 116(5), 882–892.

Wright, I. J., Reich, P. B., Westoby, M., Ackerly, D. D., Baruch, Z., Bongers, F., … others. (2004). The worldwide leaf economics spectrum. Nature, 428(6985), 821–827.

Core Team (2017). R: A language and environment for statistical computing. R Foundation for Statistical Computing, Vienna, Austria. URL https://www.R-project.org/.

